# Do AI Structure Predictors Capture Bound-State Disorder? A Benchmark on Fuzzy Protein Complexes

**DOI:** 10.64898/2026.05.30.729023

**Authors:** Juan Velasquez, Sebastien Ghent, Vladimir N. Uversky, Taseef Rahman

## Abstract

Fuzzy protein complexes, in which an intrinsically disordered protein (IDP) retains conformational disorder upon binding, pose a fundamental challenge for structure predictors trained on ordered systems, where crystal structures capture only the most ordered ensemble snapshot, making standard benchmarking metrics misleading. Here, we present the first systematic evaluation of AlphaFold3 (AF3), AlphaFold2-Multimer (AF2MM), Chai-1, and Boltz-2 on a curated dataset of fuzzy complexes from FuzDB, benchmarked against DockQ against PDB structures and NOE violation rates against manually curated BMRB restraint files, the first comprehensive collection of this kind. Across all four predictors, approximately 30% of NOE restraints were violated with nearly identical distributions regardless of predictor architecture or training data. DockQ scores fell uniformly within the Acceptable range, with AF3 marginally higher but exhibiting NOE violation rates equivalent to the weakest-performing model. Ensemble-level analysis using a first-principles implementation of the Hadzi thermodynamic model revealed that AF3 uniquely achieves near-zero mean helicity bias, in contrast to systematic overconfidence in the other predictors, yet all four models show poor per-residue helicity correlation with thermodynamic expectations. DockQ rankings reflect training data similarity to crystal structures rather than physical accuracy, and no current predictor captures fuzzy complex ensemble behavior. The *FuzzyBench-NOE* dataset, comprising NOE restraint files, predicted structures, interface hotspot annotations, and Hadzi–DSSP analysis outputs, is released on Zenodo (https://doi.org/10.5281/zenodo.20470556).

**Significance Statement:** No benchmark exists for fuzzy protein complexes, where IDPs retain disorder upon binding. We show that four state-of-the-art structure predictors violate 30% of experimental NMR distance restraints invariantly regardless of architecture, while DockQ, the standard metric, is entirely uncorrelated with this failure. Ensemble-level analysis using the Haďzi thermodynamic model reveals systematic helicity overconfidence across all predictors. Taken together, our findings imply that standard geometric metrics are fundamentally misleading for disordered systems, thus necessitating ensemble-aware evaluation.

## 1. Introduction

Intrinsically disordered proteins (IDPs) and intrinsically disordered regions (IDRs) are abundant in eukaryotic proteomes, constituting roughly 30% of eukaryotic proteins (Dunker et al., 2015), and play essential roles in signaling, transcription regulation, and the formation of membrane-less organelles (Dyson and Wright, 2005). Their disordered nature is not incidental but functional: the conformational plasticity of IDPs enables binding to multiple partners (Wright and Dyson, 2015), fine-tuning of interaction affinity in response to post-translational modifications (Bah and Forman-Kay, 2016), and context-dependent allosteric regulation unavailable to structured proteins (Berlow et al., 2018).

A significant and functionally important subset of IDPs form *fuzzy complexes*, in which structural disorder is maintained upon partner binding and is required for biological function rather than representing an incompletely resolved structure (Tompa and Fuxreiter, 2008; Fuxreiter, 2020). Fuzzy complexes exhibit a range of bound-state behaviors, including dynamic profiles, clamp conformations, structural polymorphism, and partially bound states in which the IDP engages the target with only a subset of its interface contacts (Tompa and Fuxreiter, 2008). These properties underlie important biological phenomena including allosteric coupling (Wright and Dyson, 2015), context-dependent binding specificity (Fuxreiter, 2020), and sensitivity to non-interface mutations that would be silent in ordered systems (Haďzi et al., 2021). The conformational phase space of target-bound IDPs has been formalized by Haďzi, Loris, and Lah (Haďzi et al., 2021), who showed that the bound-state ensemble is governed by two parameters derivable from sequence and structure: the IDP folding propensity (Δ*G*_CH_), computed from Lifson–Roig helix–coil theory (Lifson and Roig, 1961), and the distribution of interface interaction hotspots, characterized by the fraction of the IDP bounded by its terminal hotspots (*N*_SEP_*/N* ) and their spacing evenness (*κ_H_*). Together, these parameters define a phase diagram that predicts whether a bound IDP will be helical, disordered, adopt a clamp conformation, or populate partially bound states.

The rapid adoption of AI structure predictors following AlphaFold2 (Jumper et al., 2021) has created an urgent need to characterize their behavior on proteins that violate the assumption these models were trained on: that a protein has a single, stable structure. AlphaFold2-Multimer (Evans et al., 2021) uses paired multiple sequence alignments to capture evolutionary co-variation between interacting chains and remains widely used in practice. AlphaFold3 (Abramson et al., 2024) replaces the structure module with a diffusion-based generator, extending predictions to nucleic acids, small molecules, and modified residues. Chai-1 (Chai Discovery et al., 2024) follows a similar diffusion architecture but adds protein language model embeddings and trainable experimental constraint features. Boltz-2 (Passaro et al., 2025) is an open-source AF3-equivalent, distinguished by fully public training code and, in its latest iteration, incorporation of binding affinity prediction. On ordered complex benchmarks, all four models achieve 70–80% success under stringent criteria (Zhou et al., 2025), and a recent unified comparison via ABCFold noted that idiosyncratic strengths and weaknesses of these models remain largely uncharacterized (Elliott et al., 2024). For IDP-containing complexes specifically, AF2-Multimer achieves reasonable success on homogeneous binding modes but degrades for heterogeneous, dynamic interactions (Omidi et al., 2024), and a recent evaluation of AF3 on disorder-to-order transitions showed unpredictable multimer performance (Dao et al., 2025). Notably, AlphaFold2’s pLDDT confidence score has been shown to identify IDRs that undergo disorder-to-order transitions upon binding at up to 88% precision (Alderson et al., 2023) — a finding that is directly relevant to interpreting predictor behavior on fuzzy complexes, where retained disorder rather than induced folding is the defining feature. Sequence-based predictors of disordered binding sites, including MoRF predictors (Katuwawala et al., 2019; Song and Kurgan, 2025) and ANCHOR-based methods (Dosztanyi et al., 2009), address the complementary question of *where* an IDP engages its partner but do not predict the 3D geometry or bound-state conformational ensemble. Performance on fuzzy complexes, where the bound IDP retains dynamics by design and the correct answer is a conformational ensemble rather than a single structure, remains entirely uncharacterized. More broadly, no benchmark of any kind exists for fuzzy complex structure prediction. The field has no agreed metric, no reference dataset, and no established baseline against which predictor performance on this class of system can be measured. This absence is not accidental as fuzzy complexes resist standard benchmarking precisely because the ground truth is an ensemble rather than a structure, and assembling the experimental data necessary to define that ensemble at scale requires manual curation across heterogeneous NMR archives. We aim to fill that particular gap with the present work.

A fundamental difficulty in benchmarking predictors on fuzzy complexes is defining appropriate ground truth. For fuzzy complexes, the crystal structure represents the most ordered state of the ensemble and is therefore a biased and incomplete reference rather than the ground truth against which a predictor should be judged. Haďzi et al. (Haďzi et al., 2021) demonstrated this directly: treating the bound IDP as static overestimates mutational effects on binding affinity by orders of magnitude, because the ensemble adapts to mutations in ways a single static structure cannot represent. Existing IDP benchmarks have used crystal structures or NMR ensemble RMSD as ground truth, both of which carry this bias. NOE distance restraints from solution NMR provide an orthogonal, model-independent alternative: direct experimental measurements of inter-atom proximity in solution that do not depend on any structural reference frame, computational model, or ensemble averaging assumption.

Here we present the first systematic four-way comparison of AF3, AF2MM, Chai-1, and Boltz-2 on fuzzy protein complexes, using a total of 105 systems curated from FuzDB with manually verified BMRB NOE restraint files (Hoch et al., 2023). We evaluate each predictor using DockQ (Mirabello and Wallner, 2024) against PDB structures, NOE violation rates against experimental restraints, and a novel ensemble-level metric derived from a from-scratch implementation of the Haďzi et al. (Haďzi et al., 2021) statistical thermodynamic model, computing *p*(*w_i_*) via the full 3 × 3 Lifson–Roig transfer matrix (Lifson and Roig, 1961) with complete 2*^NH^* − 1 partial-binding subensemble enumeration, which is the physically correct treatment for fuzzy complexes in which IDPs may engage the target with any subset of their hotspot residues. This constitutes the first application of the Haďzi framework at benchmark scale across multiple predictors and a large, diverse system set. Our results reveal that DockQ rankings are artifactual, that all four predictors fail equivalently on experimental constraints, and that none captures the ensemble behavior of fuzzy complexes at the residue level.

## 2. Results

### 2.1. Dataset: 105 Fuzzy Complexes with Tiered Experimental Ground Truth

We curated a benchmark dataset of 105 IDP–target fuzzy complexes from FuzDB v4.0 (Hatos et al., 2022), requiring deposited PDB structures with experimental evidence of bound-state disorder. Systems span a range of IDP lengths (8–172 residues, median 45), interface sizes (2–51 residues, median 10), and hotspot counts (0–9, median 4), providing a diverse representation of the fuzzy complex conformational phase space (Haďzi et al., 2021). For each system, three predicted models were generated per predictor (sequence-only input, no templates), and the top three ranked models by predictor confidence were used for ensemble-level analysis.

The dataset is structured in two tiers. All 105 systems are fuzzy complexes and were evaluated using DockQ (Mirabello and Wallner, 2024) against PDB reference structures and the Haďzi–DSSP ensemble calibration metric. A subset of 75 systems additionally had usable NMR restraint files identified and manually verified in BMRB (Hoch et al., 2023), constituting the first comprehensive NOE restraint collection for fuzzy complexes, and were used for NOE violation analysis. The remaining 30 systems are also fuzzy complexes but lack deposited BMRB restraint data and were evaluated on the first two metrics only. One system (2LTO) had no interface hotspots identified by PPCheck (Sukhwal and Sowdhamini, 2015); it was retained in DockQ and NOE analyses but its Haďzi *p*(*w_i_*) profile reflects the unbound helix–coil ensemble rather than the target-bound ensemble, as no interaction weights are applied in the absence of hotspot residues.

### 2.2. DockQ: Marginal and Uniform Acceptable Performance

All four predictors produced DockQ scores predominantly in the Acceptable range (0.23– 0.49), with none reliably reaching the Medium quality threshold (*>* 0.49; Figure 1). Mean DockQ scores across all 105 systems (315 models per predictor, 312 for Chai-1) were: AF3 = 0.381 ±0.216, AF2MM = 0.364 ±0.202, Boltz-2 = 0.313 ±0.190, and Chai-1 = 0.264 ±0.188 (Table 1). AF3 achieved the highest mean DockQ, yet fewer than 4% of AF3 models reached the High quality threshold (*>* 0.80; 4/315), and 27% were scored as Incorrect (*<* 0.23; 86/315). Chai-1 had the weakest overall performance, with 49% of models classified as Incorrect (152/312) and a mean iRMSD of 7.7 ± 5.5 Å versus 5.3 ± 4.6 Å for AF3.

**Figure 1.**
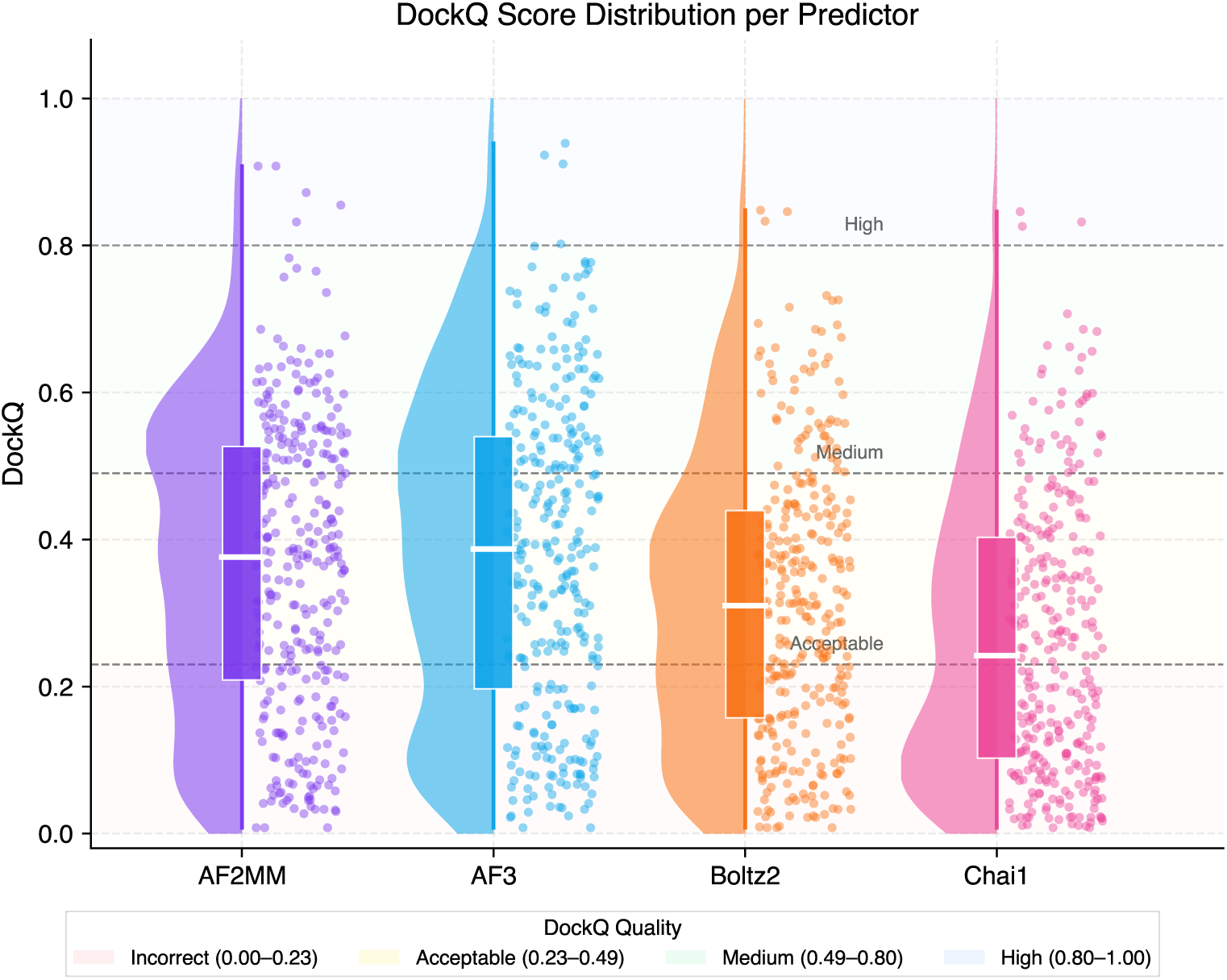
DockQ score distributions across four structure predictors on 105 fuzzy protein complexes. Each panel shows the distribution of DockQ scores for all predicted models (315 per predictor, 312 for Chai-1) as a raincloud plot combining a half-violin (left, distribution shape), boxplot (centre, median and IQR), and jittered individual data points (right). Dashed horizontal lines indicate standard DockQ quality thresholds: Acceptable (≥0.23), Medium (≥0.49), and High (≥0.80). Background shading denotes quality bands. All four predictors produce distributions centered in the Acceptable range, with no predictor consistently reaching the Medium threshold. AF3 achieves the highest median DockQ (0.387) but retains a substantial tail of Incorrect predictions (*<*0.23). Chai-1 shows the weakest overall performance, with a median of 0.242 and the largest fraction of Incorrect models (49%). The near-identical distributional shapes across architecturally distinct predictors indicate a systematic rather than implementation-specific limitation.

**Table 1:** Table I. DockQ summary across predictors (105 systems, 3 models each). Quality bins: Incorrect (Inc.) *<* 0.23, Acceptable (Acc.) 0.23–0.49, Medium/High (M/H) ≥ 0.49.

| Predictor | DockQ | Median | IQR | Fnat | iRMSD ($\text{\AA}$) | Inc. | Acc. | M/H |
| --- | --- | --- | --- | --- | --- | --- | --- | --- |
| | mean ± SD | | Q1–Q3 | mean ± SD | mean ± SD | $n$ | $n$ | $n$ |
| AF2MM | $0.364 \pm 0.202$ | 0.376 | 0.21–0.53 | $0.475 \pm 0.277$ | $5.5 \pm 4.5$ | 94 | 117 | 104 |
| AF3 | $0.381 \pm 0.216$ | 0.387 | 0.20–0.54 | $0.491 \pm 0.274$ | $5.3 \pm 4.6$ | 86 | 119 | 110 |
| Boltz-2 | $0.313 \pm 0.190$ | 0.310 | 0.16–0.44 | $0.445 \pm 0.281$ | $6.4 \pm 4.9$ | 119 | 139 | 57 |
| Chai-1 | $0.264 \pm 0.188$ | 0.242 | 0.10–0.40 | $0.372 \pm 0.283$ | $7.7 \pm 5.5$ | 152 | 117 | 43 |

Kruskal-Wallis analysis confirmed a significant overall difference in DockQ across predictors (*H* = 61.7, *p* = 5.0 × 10^−11^). However, pairwise Mann-Whitney U tests with Benjamini-Hochberg correction revealed that AF3 and AF2MM are statistically indistinguishable (*U* = 47,516, *p*_BH_ = 0.36), despite their apparent mean difference of 0.017 DockQ units. All comparisons involving Chai-1 or Boltz-2 were significant after correction (*p*_BH_ *<* 0.002). As shown below, this ranking is not recapitulated in NOE violation rates, suggesting it reflects similarity to PDB crystal structures rather than genuine differences in physical accuracy.

### 2.3. NOE Violation Rates: Invariant Failure Across All Predictors

NOE distance restraint violation analysis on the 75-system subset revealed a striking result: all four predictors violated 30–32% of experimental restraints, with essentially identical distributions regardless of predictor architecture or training data (AF2MM: 30%, AF3: 30%, Boltz-2: 30%, Chai-1: 32%; Figure 2A). The fraction of satisfied restraints was 55–56% across all models, with 6% unmapped in all cases.

**Figure 2.**
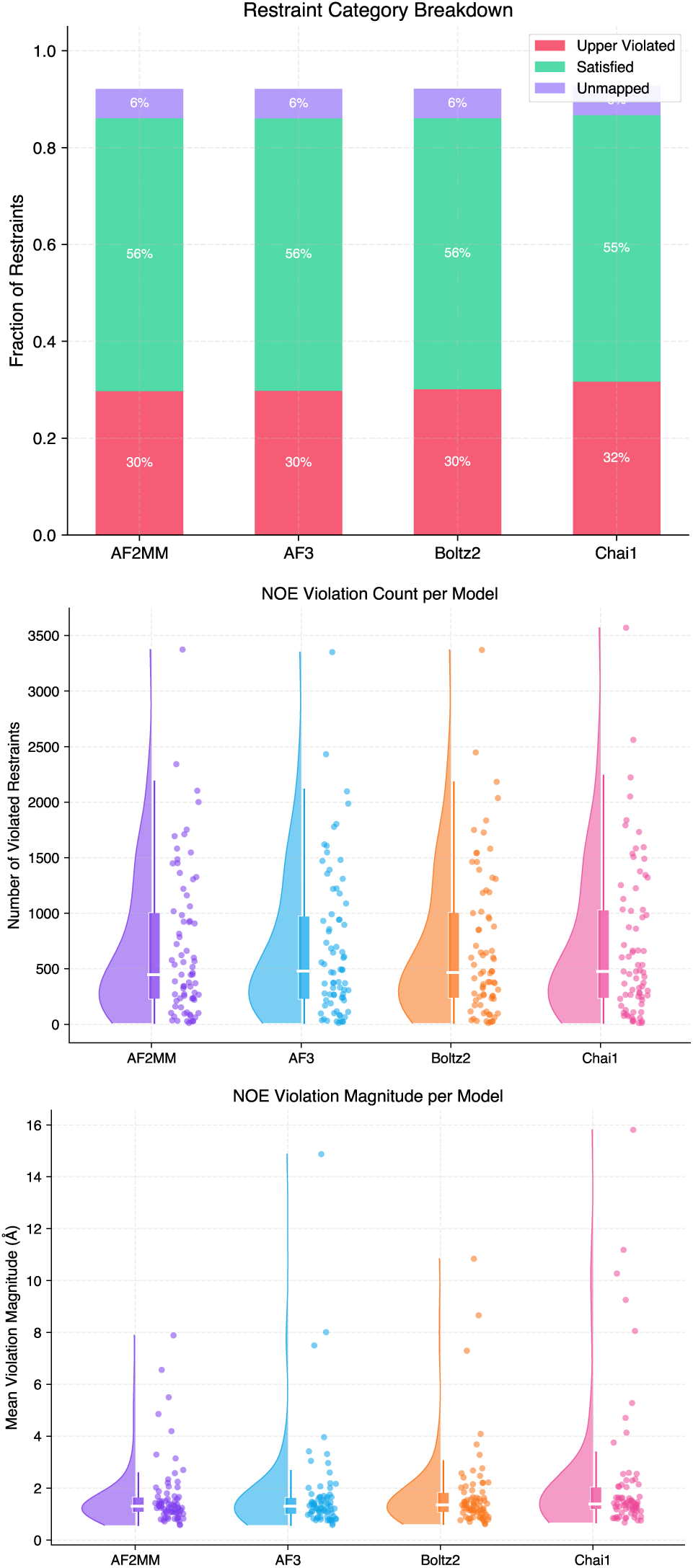
NOE distance restraint violation analysis across four structure predictors on 75 fuzzy complexes with verified BMRB restraint data. **(A)** Restraint category breakdown per predictor. All four predictors violate 30–32% of experimental NOE upperbound distance restraints (Upper Violated, red), with 55–56% satisfied (green) and 6% unmapped (purple) in all cases. The near-identical distributions across architecturally distinct models indicate a systematic rather than implementation-specific failure. **(B)** Violation count per system per predictor shown as raincloud plots (half-violin, boxplot, jittered points). Median violation counts are 450–500 restraints per system across all predictors, with highly right-skewed distributions and extreme outliers exceeding 3,000 violations, consistent with a subset of systems where predicted structures are globally incorrect. **(C)** Mean violation magnitude per system per predictor. Median magnitudes are 1.3–1.5 Å across all models, with long right-skewed tails reaching 14.9 Å for AF3 and 15.8 Å for Chai-1. The distributional shapes are indistinguishable across predictors. Together, panels A–C demonstrate that no predictor satisfies experimental NMR distance constraints more reliably than any other, despite substantial differences in DockQ performance.

Violation magnitude distributions showed median violations of approximately 1.3–1.5 Å across all predictors, with long right-skewed tails extending to 14.9 Å for AF3 and 15.8 Å for Chai-1 (Figure 2C). The interquartile ranges and overall distributional shapes were nearly identical across models. This invariance persists across two generations of prediction architecture (structure-module-based AF2MM versus diffusion-based AF3, Chai-1, and Boltz-2) and across models trained on substantially different datasets, indicating a fundamental failure mode common to the entire class of single-structure predictors when applied to fuzzy complexes. Violation counts per system were also highly variable and right-skewed, with medians around 450–500 violated restraints per system and extreme outliers exceeding 3,000 (Figure 2B).

Critically, AF3’s DockQ advantage over Chai-1 (mean 0.381 vs. 0.264, a 44% relative improvement) did not translate to any improvement in NOE satisfaction, with both models violating 30–32% of experimental restraints. This dissociation is the central empirical finding of this study: *DockQ rankings on fuzzy complexes are not predictive of experimental constraint satisfaction* and should not be used as a measure of physical accuracy for conformationally disordered systems.

### 2.4. Haďzi vs. DSSP: Ensemble Calibration Analysis

To diagnose the mechanistic basis of predictor failure, we compared per-residue DSSP helicity averaged across predicted models against the thermodynamic per-residue helical probability *p*(*w_i_*) from the Haďzi model (Table 2; Figure 3).

**Figure 3.**
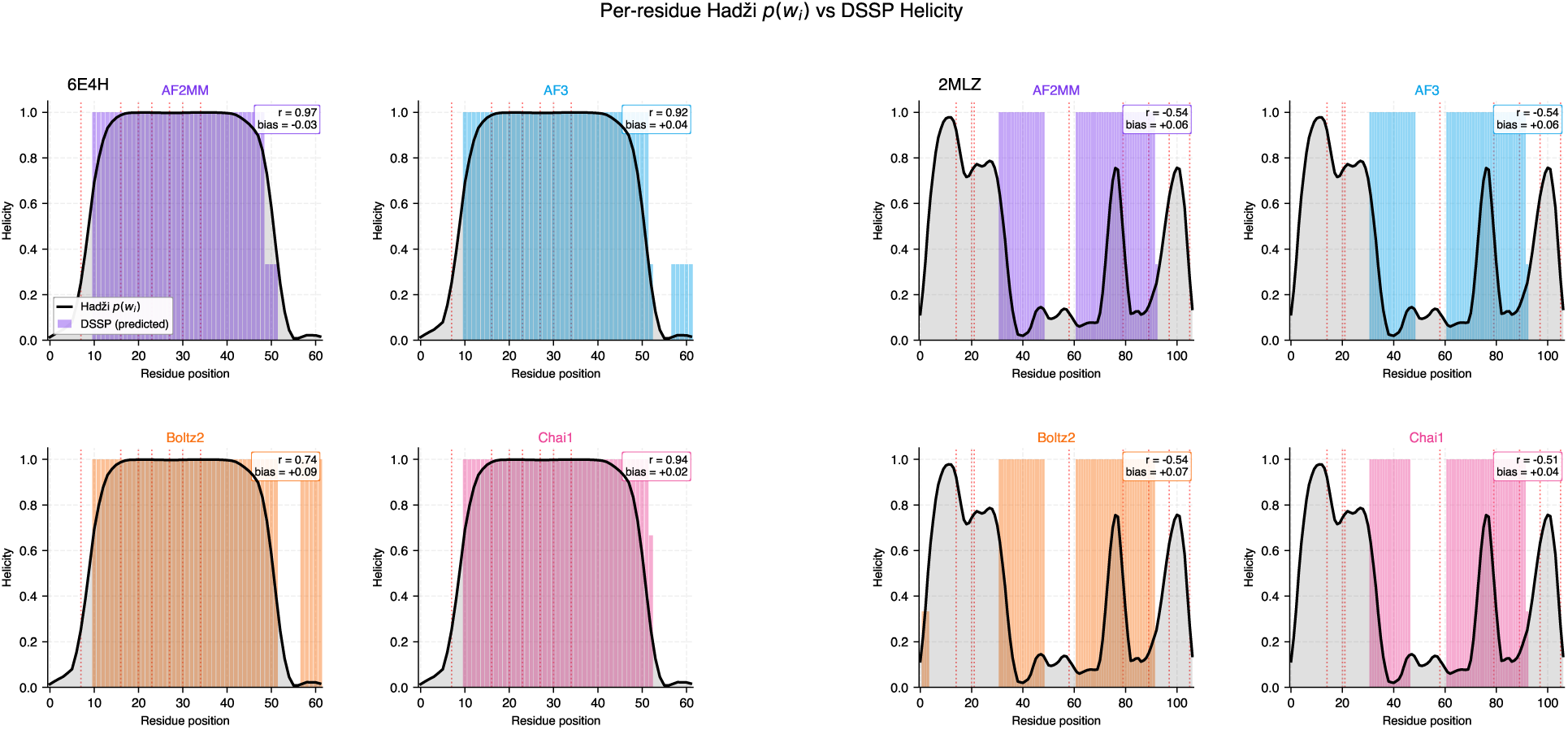
Per-residue Haďzi. *p*(*w_i_*) **vs averaged DSSP helicity for a well-predicted (6E4H) and a poorly-predicted (2MLZ) fuzzy complex.** Black lines show the thermodynamic per-residue helical probability *p*(*w_i_*) computed from the Haďzi model using each system’s IDP sequence and interface hotspot distribution. Colored bars show the per-residue DSSP helicity averaged across three predicted models per predictor. Red dashed vertical lines indicate hotspot positions. **Left (6E4H):** all four predictors track the Haďzi profile closely (Pearson *r* = 0.74–0.94), correctly capturing both the helical core and the disordered termini of this system. **Right (2MLZ):** all four predictors fail to reproduce the complex multi-modal Haďzi profile (*r* = −0.54 to −0.51), instead predicting helical blocks from residue positions 35-45 and 62-92 where the thermodynamic model expects alternating helical and disordered regions. This comparison illustrates why the population-level mean Pearson *r* ≈ 0.29 reflects genuine residue-level failures rather than statistical noise: predictors succeed on systems with simple helical profiles but collapse the conformational complexity of multi-modal fuzzy ensembles into a single ordered conformation.

**Table 2:** Table II. Haďzi–DSSP ensemble calibration analysis. Bias = mean DSSP − mean Haďzi *p*(*w_i_*). Sig. = FDR-corrected *p <* 0.05. OC/UC = severely over/underconfident (|bias| *>* 0.2).

| Predictor | Mean Bias | Pearson $r$ | Sig. | DSSP=0 | OC / UC |
| --- | --- | --- | --- | --- | --- |
| AF2MM | $+0.075 \pm 0.303$ | $0.287 \pm 0.376$ | 52/104 | 19 (18%) | 33 / 13 |
| AF3 | $-0.000 \pm 0.264$ | $0.297 \pm 0.358$ | 53/104 | 26 (25%) | 18 / 19 |
| Boltz-2 | $+0.059 \pm 0.286$ | $0.304 \pm 0.344$ | 56/104 | 17 (16%) | 29 / 15 |
| Chai-1 | $+0.075 \pm 0.304$ | $0.273 \pm 0.340$ | 52/103 | 20 (19%) | 31 / 14 |

AF3 exhibited a uniquely near-zero mean helicity bias of −0.0004 (mean DSSP helicity 0.358 vs. Haďzi expectation 0.358; Figure 4), indicating that its diffusion-based sampling produces predicted ensembles with the correct global helical fraction on average. By contrast, AF2MM, Boltz-2, and Chai-1 systematically overpredicted bound-state helicity with mean biases of +0.075 ± 0.303, +0.059 ± 0.286, and +0.075 ± 0.304 respectively. One-sample Wilcoxon signed-rank tests against zero confirmed that AF2MM and Chai-1 biases were significantly positive after Benjamini-Hochberg correction (both *p*_BH_ = 0.045), while AF3 (*p*_BH_ = 0.90) and Boltz-2 (*p*_BH_ = 0.11) were not. Critically, Kruskal-Wallis analysis showed no significant overall difference in bias distributions across predictors (*H* = 4.8, *p* = 0.19), and all pairwise comparisons were non-significant after correction (*p*_BH_ *>* 0.18), confirming that AF3’s near-zero mean bias is not statistically distinct from the positive bias of the other three models at the population level. AF3 was severely overconfident (bias *>* 0.2) in 18/104 systems, substantially fewer than AF2MM (33/104), Chai-1 (31/103), and Boltz-2 (29/104).

**Figure 4.**
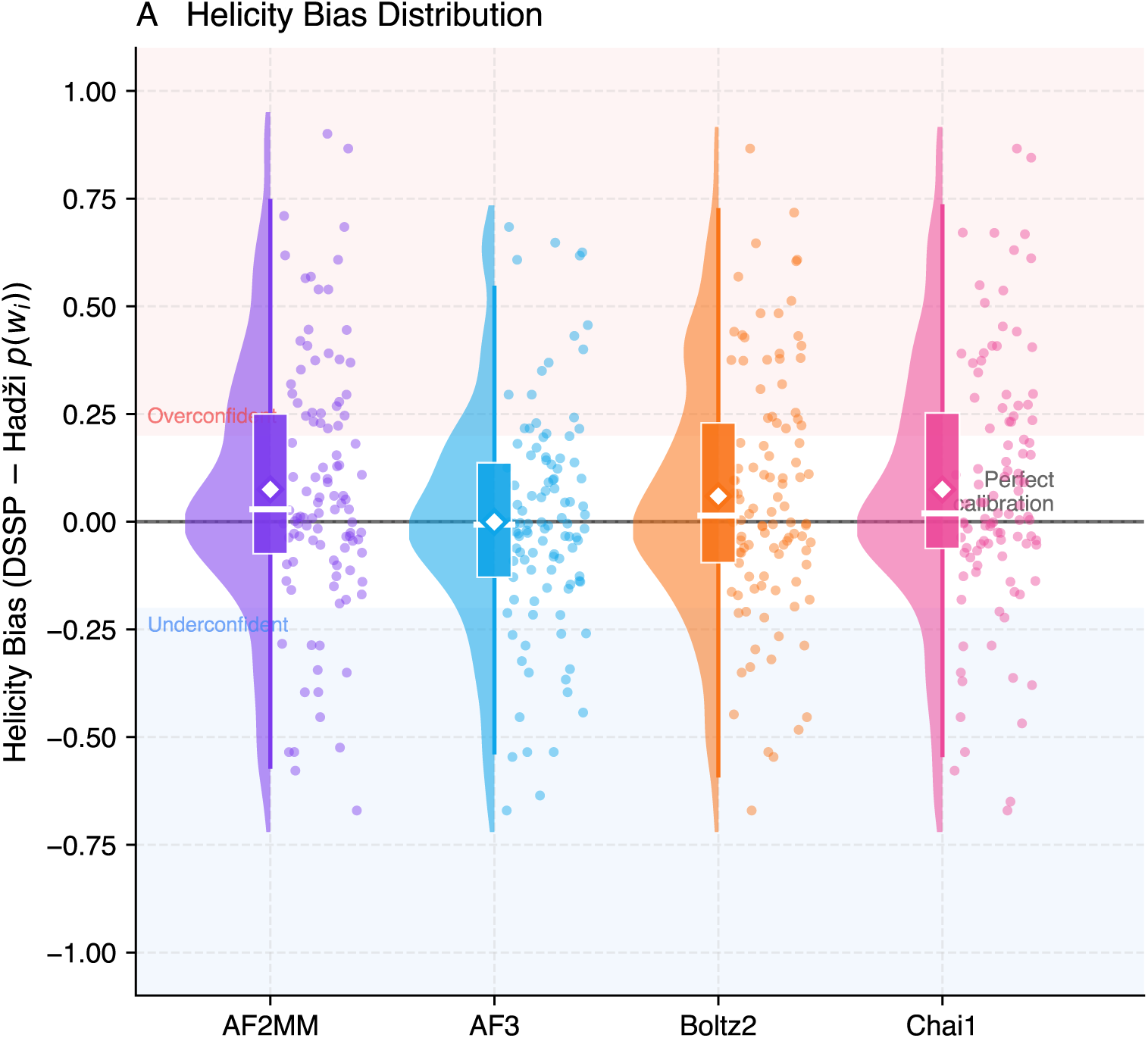
Helicity bias distributions across four structure predictors. Raincloud plots showing the distribution of per-system helicity bias (DSSP − Haďzi *p*(*w_i_*)) for each predictor across 105 fuzzy complexes. The horizontal zero line indicates perfect calibration. The red shaded region (bias *>* 0.2) denotes severely overconfident predictions, and the blue shaded region (bias *<* −0.2) denotes severely underconfident predictions. Diamond markers show the population mean per predictor. AF3 achieves a uniquely near-zero mean bias (−0.0004), indicating correct global helical fraction on average, while AF2MM, Boltz-2, and Chai-1 show systematic positive bias (+0.059 to +0.075), reflecting overconfidence in boundstate IDP helicity. However, the wide distributions and substantial tails into both shaded regions for all predictors confirm that neither systematic overconfidence nor calibration is consistent across individual systems.

However, despite AF3’s superior global calibration, per-residue correlation between DSSP and Haďzi *p*(*w_i_*) was poor across all predictors (mean Pearson *r*: AF2MM = 0.287 ± 0.376, AF3 = 0.297 ± 0.358, Boltz-2 = 0.304 ± 0.344, Chai-1 = 0.273 ± 0.340). After Benjamini–Hochberg false discovery rate correction (*α* = 0.05), approximately half of systems showed statistically significant correlations (AF2MM: 52/104, AF3: 53/104, Boltz-2: 56/104, Chai-1: 52/103), confirming that the aggregate pattern is robust and not an artifact of multiple testing. The high standard deviations in Pearson *r* reflect genuine system-to-system variability rather than noise, with some systems showing strong positive correlation and others showing near-zero or negative correlation across all predictors. Pairwise Mann-Whitney U tests confirmed that no predictor achieves significantly better residue-level correlation than any other (all *p*_BH_ *>* 0.91), demonstrating that the residue-level calibration failure is uniform across architectures.

AF3 predicted zero helicity for 25.0% of systems (26/104), compared to 18.3% for AF2MM (19/104), 19.4% for Chai-1 (20/103), and 16.3% for Boltz-2 (17/104), indicating a pronounced tendency to collapse IDP chains into completely disordered coils. This is a distinct failure mode from the systematic overconfidence of AF2MM and Chai-1: AF3 achieves correct global helicity calibration in part because its zero-helicity predictions cancel its overconfident predictions at the population level, rather than because it genuinely captures the residue-level ensemble at each system.

Taken together, these results reveal that no predictor reliably captures the per-residue helicity profile expected from the thermodynamics of the bound-state ensemble (Figure 5). AF3’s near-zero mean bias is an emergent population-level property arising from two opposing failure modes rather than genuine ensemble awareness.

**Figure 5.**
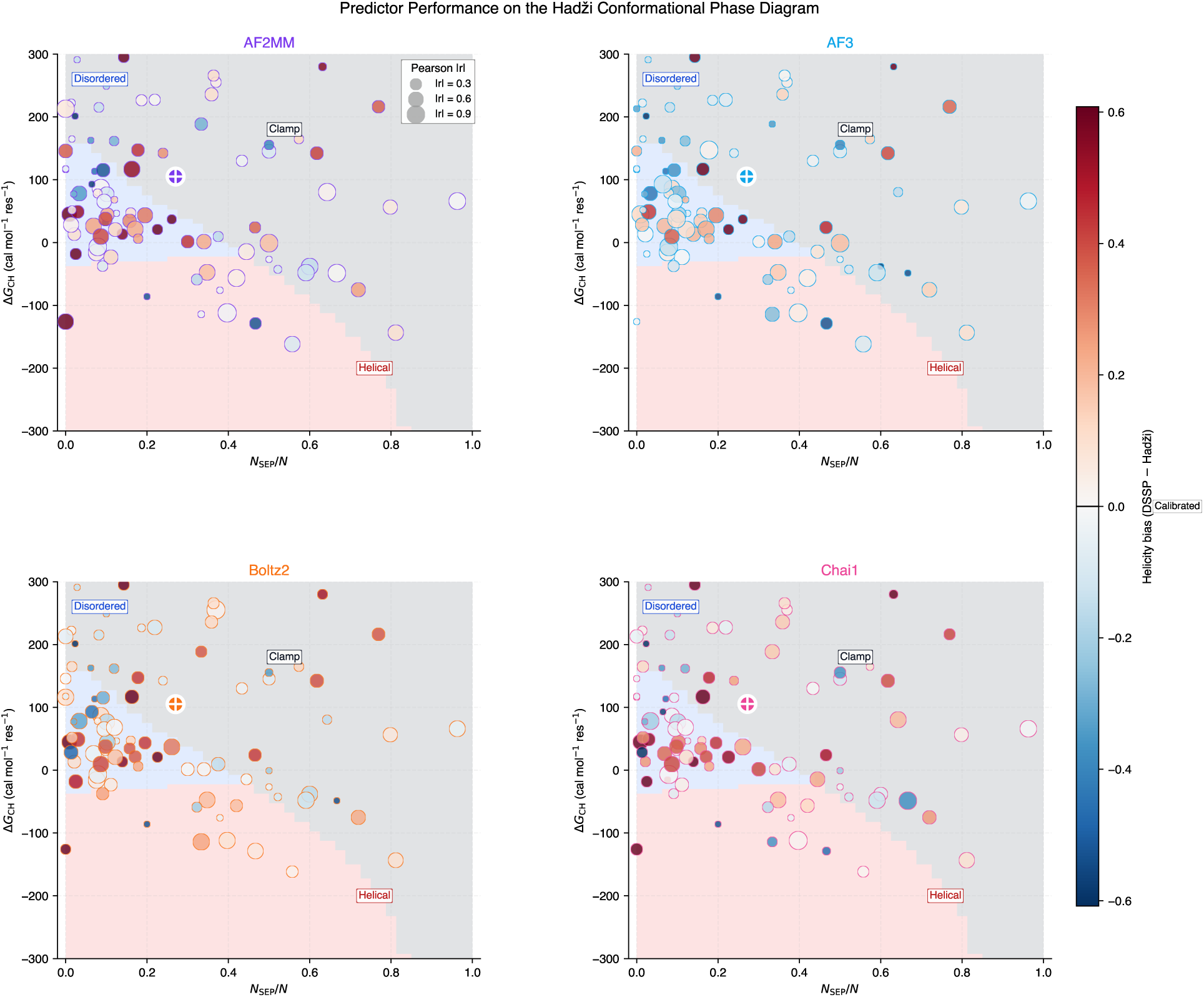
Predictor performance overlaid on the Haďzi conformational phase diagram. Each panel shows the 105 benchmark systems positioned in the thermodynamic phase space defined by *N*_SEP_*/N* (fraction of IDP bounded by terminal hotspots, x-axis) and Δ*G*_CH_ (mean IDP folding propensity, y-axis). Critically, both axes are computed from *ground truth* parameters: Δ*G*_CH_ from the real IDP sequence, and *N*_SEP_*/N* from hotspot positions identified by PPCheck on the experimental PDB crystal structure. The phase diagram background (Helical, Clamp, Disordered boundaries) is computed exactly from the Haďzi et al. (Haďzi et al., 2021) Lifson–Roig partition function using parameters matching Figure 2B of the original paper (*N* = 32, *N_H_*= 2, Δ*G*_INT_ = −3 kcal mol^−1^). The figure therefore answers: *given where each system genuinely sits in thermodynamic phase space, how well does each predictor perform there?* Point color encodes helicity bias (DSSP − Haďzi *p*(*w_i_*)): red indicates systematic overconfidence in bound-state helicity, blue indicates underconfidence, and white indicates calibrated predictions. Point size scales with |Pearson *r*|, where larger points indicate stronger residue-level correlation between predicted and thermodynamically expected helicity profiles. The crosshair marks the population mean per predictor. The benchmark dataset is concentrated in the disordered and clamp regions, confirming that the systems are genuinely fuzzy rather than simple disorder-to-order transitions. Red dots dominate the disordered region for AF2MM, Boltz-2, and Chai-1, indicating that predictors are most overconfident precisely where the ground-truth thermodynamics predicts dynamic, partially disordered bound states. AF3 shows a more heterogeneous pattern, consistent with its near-zero population-level bias arising from cancellation between overconfident and zero-helicity predictions rather than genuine ensemble awareness.

## 3. Discussion

### 3.1. Insufficiency of DockQ as a metric

This study presents the first systematic benchmark of four current-generation AI structure predictors on fuzzy protein complexes, using a dataset of 105 systems and three complementary evaluation metrics. Three independent lines of evidence converge on a unified conclusion that no current predictor captures the conformational ensemble behavior of fuzzy complexes, and standard evaluation metrics obscure rather than reveal this failure.

The 30% NOE violation rate, invariant across all four predictors, is the most direct and assumption-free demonstration of this failure. NOE restraints are direct experimental measurements of inter-atom proximity in solution and do not depend on any structural model, thermodynamic assumption, or reference frame. The observation that architecturally distinct models trained on different data produce essentially identical violation rates indicates that this is not an implementation problem solvable by better training or larger datasets within the current single-structure paradigm. A single predicted structure cannot simultaneously satisfy the full set of NOE restraints that encode a dynamic ensemble, implying that this is a category error rather than a performance gap.

The dissociation between DockQ and NOE violation rate has important implications for how the field evaluates predictors on disordered systems. AF3’s DockQ score of 0.381 is 44% higher than Chai-1’s 0.264, yet both models violate 30–32% of experimental NOE restraints. DockQ measures similarity to a crystal structure which, for fuzzy complexes, is a biased and incomplete representation of the real ensemble. The DockQ ranking mirrors the chronological order of model development rather than genuine differences in physical accuracy, consistent with the broader observation that predictor performance tracks training set coverage (Zhou et al., 2025; Omidi et al., 2024).

AF3’s near-zero helicity bias is a genuinely interesting finding that deserves mechanistic attention. The diffusion-based architecture generates structural diversity across seeds by design, which may produce a set of predicted conformations whose ensemble-average helicity fortuitously matches the thermodynamic expectation at the population level. However, the poor per-residue correlation (*r* ≈ 0.30) and the high rate of complete helicity collapse (25% of systems) reveal that this global calibration arises from two opposing failure modes cancelling rather than from genuine ensemble sampling. The per-residue profiles (Figure 3) illustrate this directly: on systems with complex multi-modal helicity profiles such as 2MLZ, all four predictors collapse the conformational heterogeneity into a single broad helical block, achieving anti-correlation with the thermodynamic expectation (*r* ≈ −0.54).

The phase diagram analysis (Figure 5) reveals that the benchmark dataset is concentrated in the disordered and clamp regions of the Haďzi phase space, confirming that the systems are genuinely fuzzy rather than borderline disorder-to-order transitions. The predominance of red points in the disordered region for AF2MM, Boltz-2, and Chai-1 demonstrates that overconfidence is specifically elevated in the phase space region where the thermodynamics predicts the most complex ensemble behavior. This is precisely where single-structure predictors are most fundamentally mismatched to the problem.

### 3.2. Relationship to pLDDT-Based Identification of Conditionally Folded IDRs

A related but distinct line of work has shown that AlphaFold2’s per-residue confidence score (pLDDT) can be used as a proxy to identify IDRs that undergo disorder-to-order transitions upon binding or post-translational modification — so-called conditionally folded IDRs (Alderson et al., 2023). At a 10% false positive rate, pLDDT achieves 88% precision in identifying such regions, a remarkable result given that conditionally folded structures were minimally represented in the AlphaFold2 training data. This finding is directly relevant to interpreting our results. Conditionally folded IDRs and fuzzy complexes represent opposite ends of the bound-state disorder spectrum: conditionally folded regions undergo genuine disorder-to-order transitions upon binding and are correctly described by a single structure, while fuzzy complexes retain dynamic disorder in the bound state and are fundamentally misrepresented by any single structure. The AF3 behavior we observe — high pLDDT with near-zero mean helicity bias at the population level — is consistent with the Alderson et al. finding: AF3 may be capturing the conditional folding signal for the subset of our fuzzy systems that are close to the helical phase boundary, while simultaneously collapsing genuinely fuzzy systems into over-confident single conformations. The 25% zero-helicity collapse rate and the anti-correlated per-residue profiles for systems such as 2MLZ (Figure 3) confirm that high pLDDT in the context of fuzzy complexes does not carry the same meaning as in conditionally folded IDRs — it reflects overconfidence rather than accurate disorder-to-order transition prediction. Together, these observations suggest that pLDDT-based confidence is informative for systems where the correct answer is a single folded structure but systematically misleading for systems where the correct answer is a dynamic ensemble.

### 3.3. Relationship to Sequence-Based Binding Site Predictors

Complementary to 3D structure prediction, a mature family of sequence-based methods predicts which residues in disordered regions are likely to engage binding partners, including Molecular Recognition Feature (MoRF) predictors and disordered binding site predictors such as ANCHOR and ANCHOR2 (Dosztányi et al., 2009), which use energy estimation to identify disordered segments that gain stability upon binding. A recent survey of 25 MoRF predictors spanning two decades of development shows that modern methods, including those leveraging protein language model embeddings such as IDBindT5 (Jahn et al., 2024), achieve balanced accuracies of ∼57% for predicting binding residues in disordered regions — competitive with alignment-based approaches (Katuwawala et al., 2019; Song and Kurgan, 2025). These sequence-based predictors address a fundamentally different question from 3D structure prediction: they identify *where* an IDP is likely to engage its partner at the residue level, without predicting the 3D geometry of the resulting complex. For fuzzy complexes specifically, MoRF and ANCHOR predictions identify the hotspot positions that are the input to the Haďzi thermodynamic model used here; in this sense, sequence-based binding site prediction and ensemble-level thermodynamic modeling are complementary rather than competing approaches. A natural extension of the present work would be to assess whether the hotspot residues identified by PPCheck on crystal structures are concordant with those predicted by sequence-based methods, and whether the conformational phase classification of a fuzzy complex can be predicted from sequence alone — a question with direct implications for large-scale proteome-wide analysis of fuzziness.

### 3.4. Limitations

Several limitations should be noted. First, the benchmark contains only fuzzy complexes, meaning we cannot directly compare predictor performance on ordered versus disordered systems from the same evaluation framework. However, we note that an ideal ordered control set is difficult to construct in practice: ordered IDP–target complexes with matched BMRB NOE restraint data are rare, size-matching to our dataset is non-trivial, and the Haďzi metric is calibrated for partial-binding physics that does not apply to fully cooperative systems. More fundamentally, the invariance of the 30% NOE violation rate across four architecturally distinct predictors — spanning two generations of methodology and trained on different data — constitutes an internal control: if the failure were predictor-specific, it would not be invariant. The finding that all models fail equivalently is itself evidence of a systematic limitation rather than a fuzziness-specific one that requires an ordered comparison to establish. Second, the Pearson *r* values (≈ 0.29) for Haďzi–DSSP correlation are low and variable across systems, making per-system interpretation unreliable; the aggregate pattern is consistent, but individual system predictions should be treated cautiously. Third, systems with zero predicted DSSP helicity (16–25% per predictor) may reflect atom-mapping failures rather than genuine predictions, and a sensitivity analysis excluding these systems is warranted. Fourth, the Haďzi model assumes cooperative binding and Lifson–Roig helix–coil parameters that may not hold universally across all fuzzy complex subtypes. Finally, the phase boundaries in Figure 5 are computed for a homopolymer IDP of length *N* = 32 with two hotspots and Δ*G*_INT_ = −3 kcal mol^−1^ as in the original Haďzi et al. figure; the positions of real systems on this diagram are approximate since they have heterogeneous sequences and variable hotspot counts.

## 4. Conclusions

We present the first benchmark of current-generation AI structure predictors on fuzzy protein complexes, using a curated dataset of 105 systems from FuzDB evaluated against DockQ, NOE violation rates from a manually assembled 75-system BMRB restraint collection, and a novel ensemble-level metric derived from a first-principles implementation of the Haďzi statistical thermodynamic model. All four predictors violate 30–32% of experimental NOE distance restraints with invariant distributions regardless of architecture or training data, demonstrating a systematic rather than implementation-specific failure. DockQ rankings are artifactual, reflecting crystal structure similarity rather than physical accuracy. AF3 uniquely achieves near-zero population-level helicity bias but does so through cancellation of opposing failure modes rather than genuine ensemble awareness. No current predictor reliably captures the residue-level conformational heterogeneity of fuzzy complexes. The 75-system *FuzzyBench-NOE* dataset — comprising NOE restraint files, predicted structures, interface hotspot annotations, and Haďzi–DSSP analysis outputs — is released as a community resource to support future development of ensemble-aware structure prediction methods.

## 5. Materials and Methods

### 5.1. Dataset Curation

Fuzzy protein complexes were curated from FuzDB v4.0 (Hatos et al., 2022), requiring deposited PDB structures with experimental evidence of bound-state disorder. IDP and target chain assignments were determined by chain order (chain A = IDP, chain B = target).

Sequences were extracted from PDB records and stored in FASTA format with IDP and target chains separated by ‘:’ for predictor input. A subset of 75 systems had usable NMR restraint files manually identified and verified in BMRB (Hoch et al., 2023), constituting the first comprehensive NOE restraint collection for fuzzy complexes.

### 5.2. Structure Prediction

Four predictors were evaluated: AlphaFold3 (AF3) (Abramson et al., 2024), AlphaFold2-Multimer v3 (AF2MM) (Evans et al., 2021), Chai-1 (Chai Discovery et al., 2024), and Boltz-2 (Passaro et al., 2025). All predictions used sequence-only input with no structural templates. Three models were generated per system per predictor, with the top three ranked by predictor confidence score selected for analysis.

### 5.3. DockQ Scoring

DockQ v2 (Mirabello and Wallner, 2024) was computed for each predicted model against the PDB reference structure using automatic chain mapping. Mean DockQ, Fnat, iRMSD, and LRMSD were computed across three models per system per predictor. Quality thresholds follow the standard DockQ classification: Incorrect (*<* 0.23), Acceptable (0.23–0.49), Medium (0.49–0.80), and High (*>* 0.80).

### 5.4. NOE Violation Analysis

BMRB restraint files were parsed to extract NOE upper distance bounds. Predicted structures were aligned to the PDB reference frame and inter-atom distances computed for each restraint atom pair. A restraint was classified as *upper violated* if the predicted distance exceeded the experimental upper bound by more than 0.5 Å (standard tolerance), *satisfied* if within the bound, and *unmapped* if the relevant atoms could not be assigned in the predicted structure. Violation counts and magnitudes were computed per system per model and aggregated across three models.

### 5.5. Hotspot Identification

Interface hotspot residues were identified using a lightweight reimplementation of the PPCheck algorithm (Sukhwal and Sowdhamini, 2015), applied to each PDB crystal structure. Residues were identified as interface contacts if their C*β* atoms (C*α* for glycine) were within 7 Å of a residue on the partner chain. Hotspots were defined as interface residues with normalized spatial residue interaction score (NESRI *>* 1) and normalized energy contribution (NEEC *>* 1), following the PPCheck protocol. One system (2LTO) yielded no hotspot residues and was excluded from the Haďzi–DSSP ensemble analysis but retained in DockQ and NOE analyses.

### 5.6. Haďzi Ensemble Analysis

Per-residue helical probabilities *p*(*w_i_*) were computed by a from-scratch implementation of the Haďzi et al. (2021) (Haďzi et al., 2021) statistical thermodynamic model, verified against the reference code (github.com/sanHadzi/PNAS_fuzzy). The implementation uses the full 3 × 3 Lifson–Roig transfer matrix (Lifson and Roig, 1961):

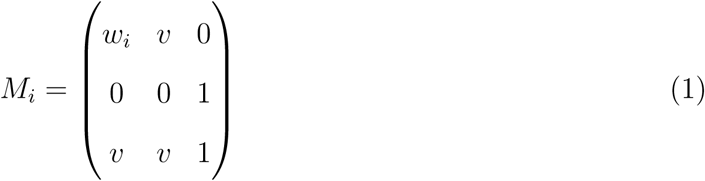

where *w_i_* is the per-residue helix propagation constant (Chakrabartty et al., 1994) pre-scaled by 2.4 to reflect the buried IDP–target interface environment. The nucleation constant *v* = 0.048 was applied uniformly. For hotspot residues, *w_i_* is multiplied by the interaction weight *x*, where Δ*G*_INT_ = −*RT* ln *x* (default −1.2 kcal mol^−1^ hotspot^−1^). The bound-state partition function *Z_B_* was computed using the complete combinatorial subensemble enumeration (all 2*^NH^* − 1 non-empty hotspot subsets, Eq. 7 of Haďzi et al.), correctly representing the partial-binding physics of fuzzy complexes. Per-residue helical probability was computed as *p*(*w_i_*) = *Z*^−1^ · *dZ_B_/d* ln *w_i_* (Eq. 9). DSSP was applied to each predicted model, with secondary structure assignments converted to binary helicity (1 = helix, 0 = other) and averaged across three models. Pearson and Spearman correlations between *p*(*w_i_*) and averaged DSSP profiles were computed per system per predictor. Pearson *p*-values were corrected using the Benjamini–Hochberg false discovery rate procedure (Benjamini and Hochberg, 1995) at FDR = 0.05. Helicity bias was defined as DSSP − *p*(*w_i_*).

### 5.7. Phase Diagram Computation

The Haďzi conformational phase diagram (Figure 5) was reproduced exactly using the Lifson– Roig partition function with parameters matching Figure 2B of Haďzi et al.: homopolymer IDP of length *N* = 32, two hotspots, Δ*G*_INT_ = −3 kcal mol^−1^. For each point on an 80 × 80 grid of (*N*_SEP_*/N* , Δ*G*_CH_) values, ⟨*n_w_*⟩ was computed via finite difference of ln *Z_B_* and the ensemble classified as: helical (⟨*n_w_*⟩ ≥ *N/*2 − 1 and ⟨*n_w_*⟩ ≥ *N*_SEP_ + 1), clamp (⟨*n_w_*⟩ *< N*_SEP_ + 1), or disordered (otherwise).

### 5.8. Statistical Analysis

All statistical tests were performed without external libraries using pure Python implementations. Pairwise differences in DockQ, helicity bias, and Pearson *r* between predictors were assessed using two-sided Mann-Whitney U tests with normal approximation. An omnibus Kruskal-Wallis test was applied before pairwise comparisons for each metric. Systematic deviation of helicity bias from zero was assessed using one-sample Wilcoxon signed-rank tests. NOE violation rate differences between predictors were assessed using Fisher’s exact test on aggregate violation counts. All *p*-values were corrected for multiple comparisons using the Benjamini-Hochberg false discovery rate procedure (Benjamini and Hochberg, 1995) at *α* = 0.05. Statistical significance thresholds: ^∗^*p*_BH_ *<* 0.05, ^∗∗^*p*_BH_ *<* 0.01, ^∗∗∗^*p*_BH_ *<* 0.001.

### 5.9. Data and Code Availability

The *FuzzyBench-NOE* dataset, including NOE restraint files, predicted structures, interface hotspot annotations, and Haďzi–DSSP analysis outputs, is deposited on Zenodo (https://doi.org/10.5281/zenodo.20470556).

## Author Contributions

Juan Velasquez: Data curation, Formal analysis. Sebastien Ghent: Data curation, Formal analysis. Vladimir N. Uversky: Writing – review and editing, Validation. Taseef Rahman: Conceptualization, Methodology, Software, Supervision, Writing – original draft.

## Acknowledgements

This project was supported by the Bellini College of Artificial Intelligence, Cybersecurity, and Computing, the Florida High Tech Corridor’s Undergraduate Research Initiative, and the CRA UR2PhD Program.

## Artificial Intelligence Disclosure

The authors used Claude (Anthropic) as an AI writing assistant to assist with editing and restructuring of manuscript text. All scientific content, interpretations, and conclusions were reviewed and verified by the authors, who take full responsibility for the accuracy and integrity of all content.

